# Human papillomavirus integration transforms chromatin to drive oncogenesis

**DOI:** 10.1101/2020.02.12.942755

**Authors:** Mehran Karimzadeh, Christopher Arlidge, Ariana Rostami, Mathieu Lupien, Scott V. Bratman, Michael M. Hoffman

**Affiliations:** Department of Medical Biophysics, University of Toronto, Toronto, ON, Canada; Princess Margaret Cancer Centre, Toronto, ON, Canada; Vector Institute, Toronto, ON, Canada; Department of Computer Science, University of Toronto, Toronto, ON, Canada

## Abstract

Human papillomavirus (HPV) drives almost all cervical cancers and up to ∼70% of head and neck cancers. Frequent integration into the host genome occurs only for tumourigenic strains of HPV. We hypothesized that changes in the epigenome and transcriptome contribute to the tumourigenicity of HPV. We found that viral integration events often occurred along with changes in chromatin state and expression of genes near the integration site. We investigated whether introduction of new transcription factor binding sites due to HPV integration could invoke these changes. Some regions within the HPV genome, particularly the position of a conserved CTCF binding site, showed enriched chromatin accessibility signal. ChIP-seq revealed that the conserved CTCF binding site within the HPV genome bound CTCF in 4 HPV^+^ cancer cell lines. Significant changes in CTCF binding pattern and increases in chromatin accessibility occurred exclusively within 100 kbp of HPV integration sites. The chromatin changes co-occurred with out-sized changes in transcription and alternative splicing of local genes. We analyzed the essentiality of genes upregulated around HPV integration sites of The Cancer Genome Atlas (TCGA) HPV^+^ tumours. HPV integration upregulated genes which had significantly higher essentiality scores compared to randomly selected upregulated genes from the same tumours. Our results suggest that introduction of a new CTCF binding site due to HPV integration reorganizes chromatin and upregulates genes essential for tumour viability in some HPV^+^ tumours. These findings emphasize a newly recognized role of HPV integration in oncogenesis.

## 1 Introduction

HPVs induce epithelial lesions ranging from warts to metastatic tumours^1^. Of the more than 200 characterized HPV strains^2^, most share a common gene architecture^3^. As the most well-recognized HPV oncoproteins, E6 and E7 are essential for tumourigenesis in some HPV^+^ tumour models^4,5,6^.

Beyond the oncogenic pathways driven by E6 and E7, emerging evidence suggests that high-risk HPV strains play an important role in epigenomic regulation of tumourigenesis. While benign papillomas usually have episomal HPV^3^, over 80% of HPV^+^ invasive cancers have integrated forms of HPV. Several studies indicate dysregulation of the transcriptome and epigenome upon integration^7,8,9^. Our knowledge of the mechanism and impact of this dysregulation, however, remains quite limited.

High-risk HPV strains have a conserved binding site for the CTCF transcription factor^10^. CTCF binds to the episomal (circular and non-integrated) HPV at the position of this sequence motif and regulates the expression of E6 and E7^10^. CTCF and YY1 interact by forming a loop which represses the expression of E6 and E7 in episomal HPV^11^. HPV integration may disrupt this loop and thereby lead to upregulated E6 and E7.

CTCF has well-established roles in regulating the 3D conformation of the human genome^12^. CTCF binding sites mark the boundaries of topological domains by blocking loop extrusion through the cohesin complex^13^. Mutations disrupting CTCF binding sites reorganize chromatin, potentially enabling tumourigenesis^14,15,16^.

Introduction of a new CTCF binding site by HPV integration could have oncogenic reverberations beyond the transcription of E6 and E7, by affecting chromatin organization. Here, we investigate this scenario—examining how HPV integration in tumours results in local changes in the epigenome, gene expression, and alternative splicing—and propose new pathways to tumourigenesis driven by these changes.

## 2 Results

### 2.1 CTCF binds a conserved binding site in the host-integrated HPV

#### 2.1.1 A specific CTCF sequence motif occurs more frequently in tumourigenic HPV strains than any other motif

We searched tumourigenic HPV strains’ genomes for conserved transcription factor sequence motifs. Specifically, we examined 17 HPV strains in TCGA head and neck squamous cell carcinoma (HNSC)^17^ and cervical squamous cell carcinoma (CESC)^18^ datasets^19^. In each strain’s genome, we calculated the enrichment of 518 JASPAR^20^ transcription factor motifs (Figure 1a). ZNF263 and CTCF motifs had significant enrichment at the same genomic regions within several tumourigenic strains (*q* < 0.05). Only in CTCF motifs, however, did motif score enrichment in tumourigenic strains exceed that of non-tumourigenic strains (two-sample t-test *p* = 0.02 *t* = −2.2). The CTCF sequence motif at position 2,916 of HPV16 occurred in the highest number of HPV strains (10/17 strains) compared to any other sequence motif (Figure 1a). This position also overlapped with the strongest CTCF ChIP-seq signal observed in the uterine squamous cell carcinoma cell line SiHa^21^ (Figure 1a). The HPV16 match’s sequence TGGC**A**CCAC**T**TGGTGGTTA closely resembled the consensus CTCF binding sequence^20^, excepting two nucleotides written in bold (*p* = 0.00001; *q* = 0.21).

**Figure 1:**
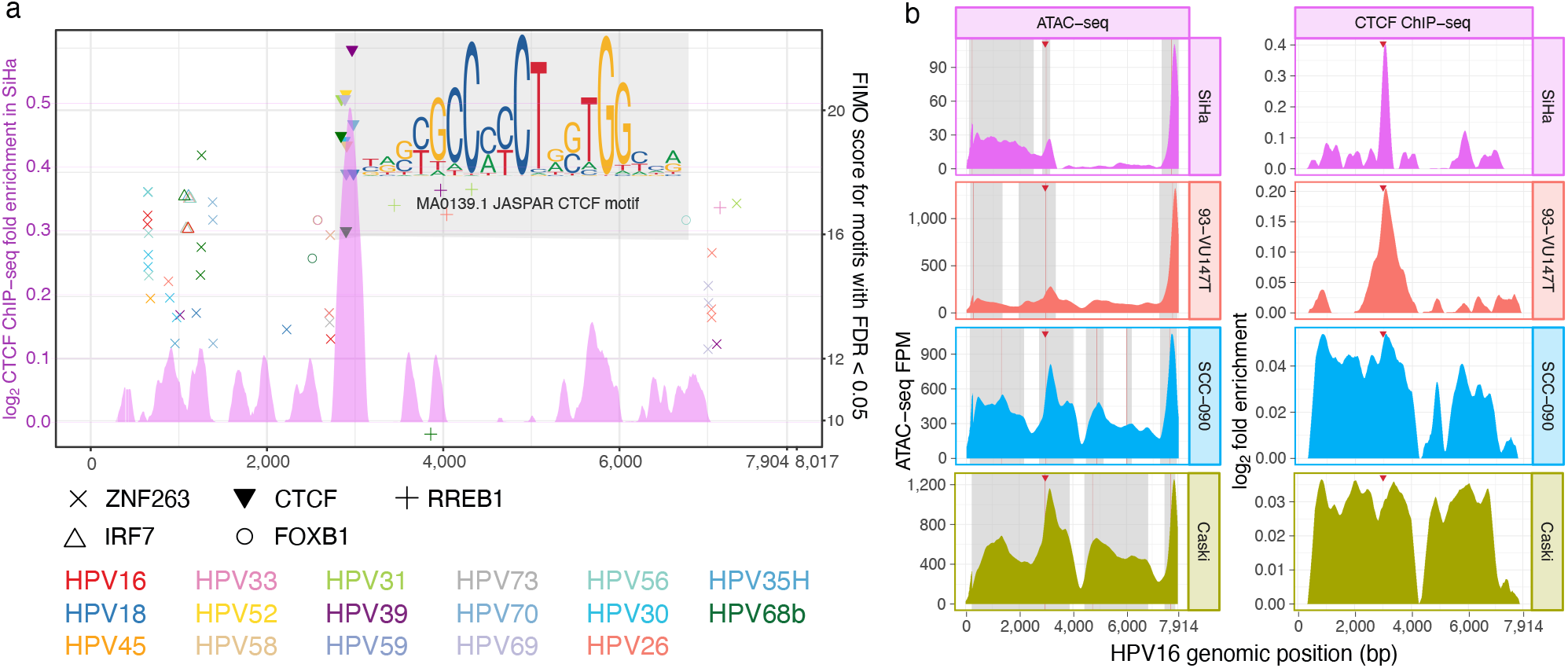
CTCF binds to its conserved binding site in HPV. **(a)** Chromatin accessibility and transcription factor motif enrichment within the HPV genome. Horizontal axis: HPV genomic position (7904 bp for HPV16 and 8017 bp for the longest HPV genome among the 17). Pink signal: CTCF MACS2 log_2_ fold enrichment over control within the HPV16 genome in SiHa. Points: FIMO^22^ enrichment scores of sequence motif matches (*q* < 0.05) of motifs occurring in at least 2/17 tumourigenic strains; symbols: motifs; colors: HPV strains. Gray area: all shown matches for the CTCF motif and its sequence logo^23^. We showed the logo for the reverse complement of the JASPAR^20^ CTCF motif (MA0139.1) to emphasize the `CCCTC` consensus sequence. **(b)** ATAC-seq MACS2 FPM (left) and CTCF ChIP-seq MACS2 log_2_ fold enrichment over the HPV16 genome (right) for 4 cell lines. To indicate no binding for regions with negative CTCF ChIP-seq log_2_ fold enrichment signal, we showed them as 0. Gray panels: chromatin accessibility peak. Red vertical line: summit of chromatin accessibility peak. Red triangle: position of the conserved CTCF sequence motif in HPV16. Dashed lines: HPV integration sites in each of the 5 cell lines 93-VU147T (orange), Caski (moss), SCC-090 (blue), and SiHa (pink).

#### 2.1.2 CTCF binds its conserved binding site in host-integrated HPV16

To test the function of the conserved CTCF motif in host-integrated HPV16, we performed ATAC-seq, CTCF ChIP-seq, and RNA-seq on 5 HPV16^+^ cell lines: 93-VU147T^24^ (7 integration sites), Caski^25^ (6 integration sites), HMS-001^26^ (1 integration site), SCC-090^27^ (1 integration site), and SiHa^21^ (2 integration sites). Unlike the other 4 cell lines, HMS-001 has only one incomplete integration of HPV into the host genome, and it lacks the genomic region containing the conserved CTCF motif^26^. For this reason, we only used HMS-001 within the comparison group. In each of the other 4 cell lines, the strongest CTCF ChIP-seq peak of the HPV genome aligned to the conserved CTCF sequence motif described above (Figure 1b, right). In each of the 4 cell lines, the second-strongest chromatin accessibility peak aligned to both the CTCF sequence motif and the CTCF ChIP-seq peak (Figure 1b, left).

The presence of both episomal and host-integrated HPV complicates the interpretation of HPV genomic signals. SiHa, however, does not contain episomal HPV^28^,29. All of the ATAC-seq and RNA-seq SiHa fragments mapping to the integration site close to the conserved CTCF motif (HPV16:3,131), also partially mapped to chr13:73,503,424. This also occurred for 3 of the 21 unpaired CTCF ChIP-seq reads mapping to HPV16:3,131. In agreement with previous reports^28^,29, these results suggest that the SiHa signal comes from the host-integrated HPV and that CTCF binding persists after HPV integration.

### 2.2 HPV integration dysregulates chromatin accessibility and transcription

#### 2.2.1 HPV dysregulates the local chromatin and transcriptome of a TCGA tumour

Integration of HPV into the host genome generates *chimeric* sequences which partially map to the host genome and partially map to the virus genome. We characterized high-confidence HPV integration sites containing chimeric sequences from TCGA cases (subsection 4.9; Supplementary Table 1). Of the TCGA HNSC patients, 9 have matched RNA-seq data measured in reads per million mapped reads (RPM) and ATAC-seq data measured in fragments per million (FPM)^30^. Using the RNA-seq data, we identified an HPV integration site in TCGA-BA-A4IH at chr9:99,952,156. The transcriptome and chromatin accessibility of this patient differed greatly from the other 8 patients at the HPV integration site (Figure 2). The other 8 patients lacked transcription (RPM < 1) or chromatin accessibility (FPM < 0.2) within 5 kbp of the integration site. TCGA-BA-A4IH, however, exhibited both active transcription and open chromatin (Figure 2a). In fact, TCGA-BA-A4IH’s chromatin accessibility and RNA expression exceeded the other 8 patients up to 400 kbp beyond the integration site (Figure 2b). Within those bounds, TCGA-BA-A4IH’s chromatin accessibility peaks often had signal exceeding that of all 8 other patients (Figure 2c).

**Figure 2:**
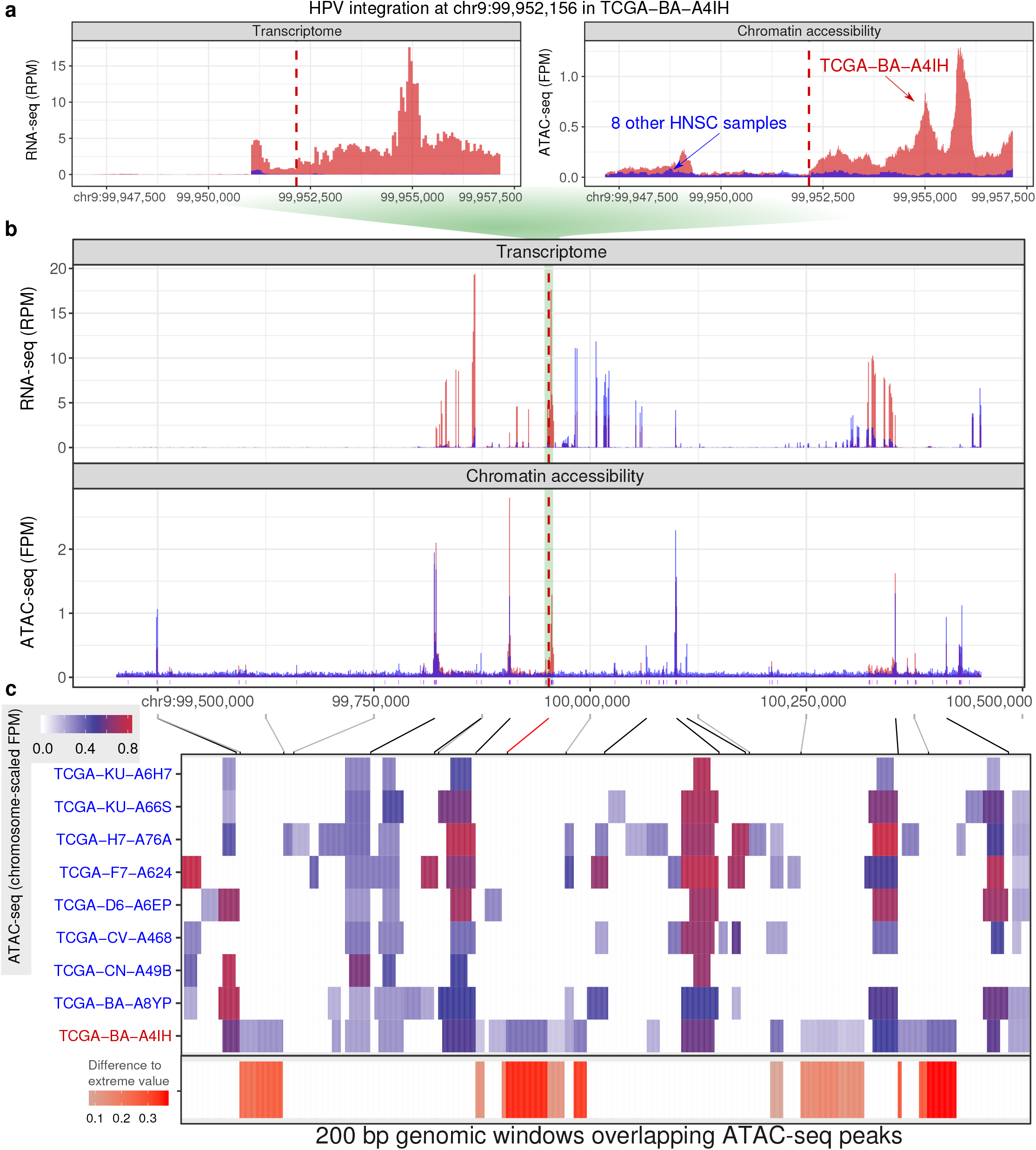
HPV integration alters the local transcriptome and epigenome. **(a)** A 10 kbp genomic window centered on TCGA-BA-A4IH’s HPV integration site. RPM RNA expression (left); FPM chromatin accessibility (right). Red: signal from TCGA-BA-A4IH; blue: signal from each of the 8 other HNSC samples. Vertical dashed red line: integration site. **(b)** Same data as (a), but in an expanded 1 Mbp genomic window. The green background shows how the coordinates of (a) fit in (b). The purple vertical bars show position of all ATAC-seq peaks found in any of the 9 tumour samples. **(c)** *(Top)*: Mapping of genomic positions for peaks with outlier signal in TCGA-BA-A4IH (gray), the position of the HPV integration site (red), and each 250,000 bp tick mark to ATAC-seq peaks. Gray diagonal lines map each 250,000 bp to the corresponding peaks. The black lines map the genomic position of the top 9 peaks with the strongest FPM in any of the 9 samples to the corresponding peaks. *(Middle)*: Heatmap of ATAC-seq peaks in the same 1 Mbp genomic window. Colour indicates ATAC-seq FPM divided by the maximum FPM value of chr9 in each patient (see subsubsection 4.5.2). Each column shows a 200 bp genomic window overlapping a peak in any of the 9 patients. We showed all 200 bp genomic windows with sliding windows of 50 bp if the window overlaps a peak. *(Bottom)*: Difference of the values in TCGA-BA-A4IH and the most extreme value in the other 8 patients when TCGA-BA-A4IH had the most extreme value among the 9 patients. We used white when TCGA-BA-A4IH did not have the most extreme value.

#### 2.2.2 HPV dysregulates local chromatin and transcriptome in HPV^+^ cell lines

To investigate the generalizability of dysregulated chromatin and transcriptome in TCGA-BA-A4IH, we conducted a similar analysis on 5 HPV^+^ lines. For each HPV integration site, we compared the cell line with integrated HPV to the other 4 cell lines without HPV at that genomic position. Only the cell line with HPV integration displayed strong expression of nearby genes (Figure 3a, top).

**Figure 3:**
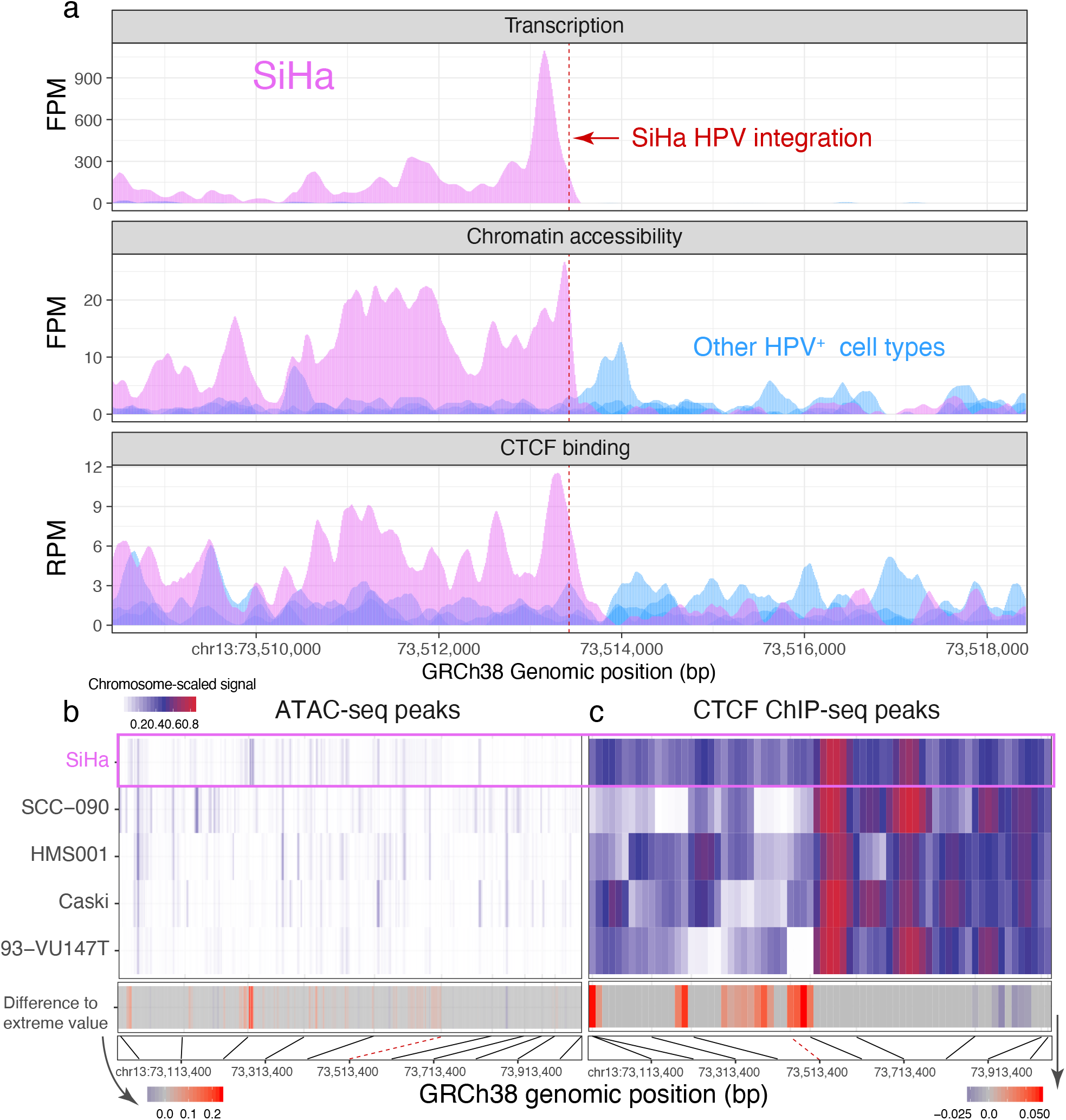
HPV integration disrupts local host epigenome and transcriptome. **(a)** Genomic assay signal for an HPV integration site of SiHa (chr13:73,513,424). Top: RNA expression FPM; middle: ATAC-seq FPM; bottom: CTCF ChIP-seq RPM. Pink bars: signal from SiHa; blue bars: signal from 4 other HPV^+^ cell lines without integration at this position. Red dashed line: HPV integration site. **(b)** ATAC-seq peaks in a 1 Mbp window centered on SiHa’s integration site. Each column shows a 200 bp genomic window overlapping a peak. We generated all 200 bp genomic windows with a stride of 50 bp which overlapped a peak in any of the 5 cell lines. *(Top)*: ATAC-seq FPM and CTCF ChIP-seq log_2_ fold enrichment over control for each cell line divided by the cell line’s corresponding maximum value in chromosome 13. *(Middle)*: Difference in the epigenome of SiHa and the most extreme value in the other 4 cell lines when SiHa had the most extreme value among the cell lines. When SiHa did not have the most extreme value, we used white. *(Bottom)*: Physical location of peaks. Black lines map every 100 kbp to the corresponding peak. Red dashed line: HPV integration site. **(c)** Similar to (b), but for CTCF ChIP-seq instead of ATAC-seq.

For each viral integration site, expression of the chimeric transcript occurred either only upstream (for 3 of the integration sites of 93-VU147T and 2 of the integration sites of Caski) or only downstream (the other 12 integration sites), never in both directions (Figure 2a). Directional chimeric transcription suggests that only one end of the integrated virus drove expression that continued past the integration site into the host genome.

Since we identified the integration site by detecting chimeric transcripts in RNA-seq data, we expected to observe transcription of the host genome at the site of viral integration. Nevertheless, transcription of these regions necessitates an active viral-dependent mechanism, as they are not transcribed in cell types without HPV integration in the same genomic regions (Figure 3a, top). Among all HPV integration sites, expression of the viral-host chimera co-occurred with chromatin accessibility signal (Figure 3a, middle). The overlap of transcription and chromatin accessibility suggests that viral integration introduces *cis*-regulatory elements which actively transcribe the viral-host chimera. The consistent recruitment of CTCF at HPV integration sites in 4 different cell lines and altered CTCF binding around integration sites suggest that CTCF plays a role in integration-dependent HPV tumourigenesis (Figure 3a, bottom).

To understand the spatial effect of HPV integration on chromatin, we examined CTCF ChIP-seq and chromatin accessibility peaks in SiHa within 500 kbp of its chr13:73,513,424 integration site (Figure 3b–c). Some of the regions of inaccessible chromatin in 93-VU147T, Caski, and SCC-090 are accessible in SiHa within 500 kbp of this integration site. In many of these regions, SiHa had more accessible chromatin compared to any of the other 4 cell lines (Figure 3b, middle). For CTCF, however, some genomic regions showed enrichment and other genomic regions showed depletion in CTCF binding (Figure 3c).

### 2.3 Integration of HPV dysregulates expression and alternative splicing of local genes

#### 2.3.1 HPV integration alters gene expression in HPV^+^ cell types

To determine whether HPV integration significantly changed gene expression, we examined changes in transcription of individual genes, as measured in transcripts per million (TPM). We used two criteria to identify outlier changes in gene expression which occurred due to HPV integration. First, we calculated expression fold change dividing log_2_ TPM in the sample with HPV integrated at some locus (TPM_HPV_+) by median TPM in samples without HPV integrated at that locus (⟨ TPM_other_ ⟩). For an HPV^+^ cell line, we only considered a gene an outlier if its expression fold change exceeded 2 (see subsubsection 4.3.3).

Out of the 17 HPV integration sites, 10 had upregulated genes only (expression fold change > 2), 3 had downregulated genes only (expression fold change < −2), and 1 (chr17:38,267,231 of 93-VU147T) had both upregulated and downregulated genes (Figure 4a, middle).

**Figure 4:**
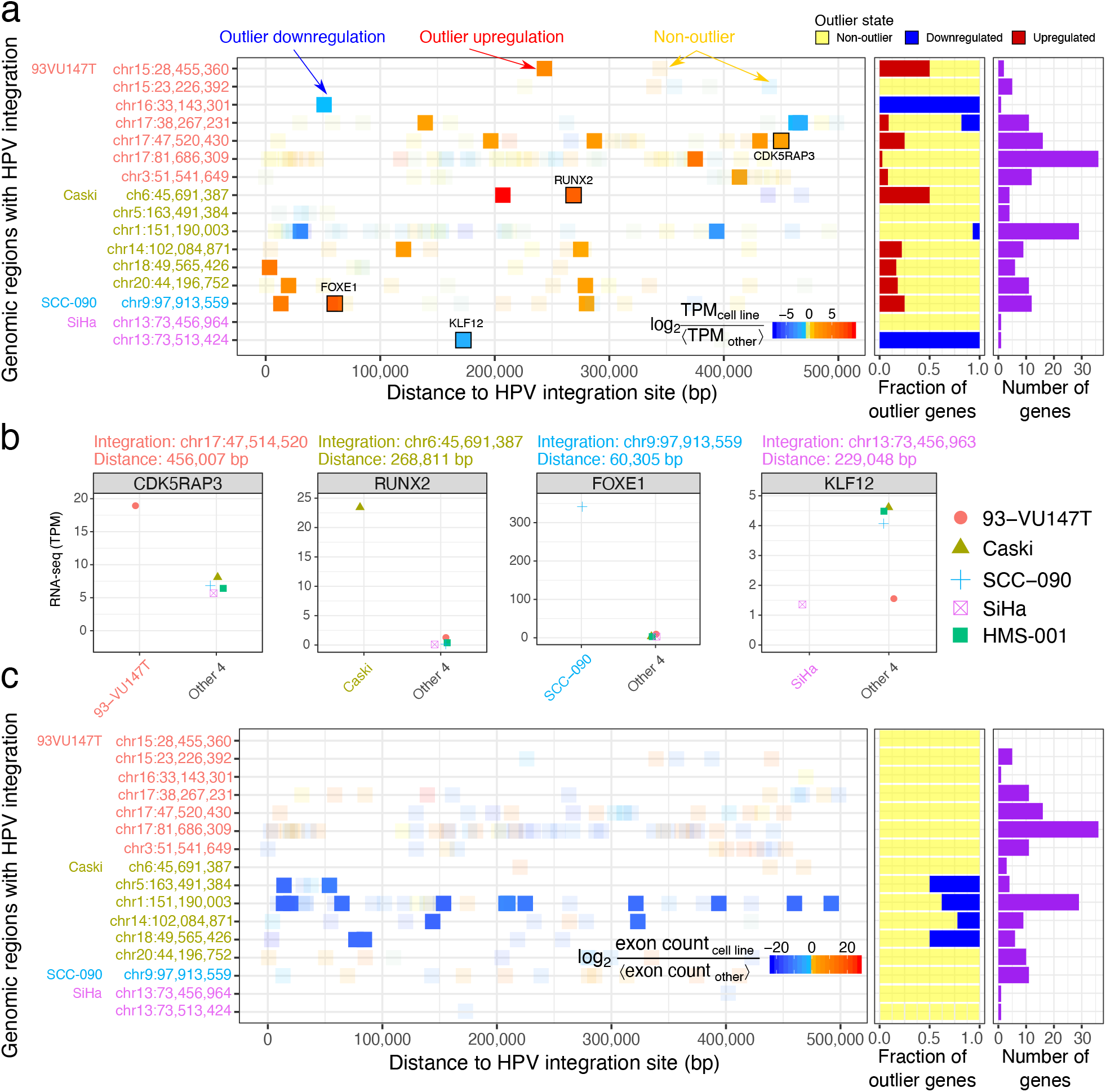
HPV integration alters local transcription and splicing. **(a)** (*Left*) Distances between 166 Ref-Seq genes within 500 kbp of 17 HPV integration sites. Colour: log_2_ TPM of the cell line with HPV integration divided by median TPM of the other 4 cell lines. Solid squares: 20 upregulated (red) and 4 downregulated (blue) outlier genes. Transparent squares: 140 genes without outlier change in gene expression. (*Middle*): Fraction of genes within 500 kbp of each HPV integration site which are either non-outlier (yellow), downregulated (blue), or upregulated (red). We labeled one gene from each cell type and visualized their TPM in (b). (*Right*): Number of genes within 500 kbp of each HPV integration site. For the overlapping integration sites in SiHa, we showed each gene in only one row to avoid duplication. **(b)** Expression of one outlier gene from each of the 5 cell lines compared to the other 4 cell lines without HPV around the gene. **(c)** Similar to (a) but for differential exon usage of 159 Ensembl genes within 500 kbp of 17 HPV integration sites. Colour: DEXSeq model fold change in exon count for the exon with the most extreme change in expression. Solid squares: 19 genes with DEXSeq *q* < 0.2 and absolute exon fold change > 1. Transparent squares: 140 genes without outlier change in exon usage.

#### 2.3.2 HPV integration sites alter gene splicing

Our results suggested that HPV integration increases chromatin accessibility and alters CTCF binding. Since chromatin-binding proteins, including CTCF, can modify gene splicing^31^, we investigated whether HPV integration affects alternative splicing of nearby genes.

We quantified how the expression of each exon varies independent of the global expression of that gene (see subsubsection 4.3.2). HPV integration sites in Caski and SCC-090 displayed outlier expression of specific exons of genes within 500 kbp (Figure 4c). These results indicate that HPV integration can influence differential exon usage of neighbouring genes.

### 2.4 HPV modifies the epigenome and transcriptome within 100 kbp of integration sites

The dysregulation of gene expression and splicing near HPV integration sites may relate to altered chromatin structure. We investigated transcriptomic and epigenomic dysregulation upon HPV integration in the RNA-seq, ATAC-seq, and CTCF ChIP-seq data. At each integration site, we compared the genomic coverage of each assay for the cell line with HPV integration to the average in the other four cell lines:

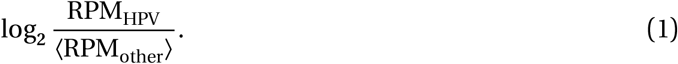

This allowed us to distinguish sample-specific variability from variations due to HPV integration.

We calculated RPM fold change (Equation 1) for all 10 kbp genomic windows around any HPV integration site. We calculated the same measurement for 10 random permutations of HPV integration sites. For each permutation, we moved the location of each HPV integration site in each cell line to a random integration site from another cell line, without replacement. We scrambled only the locations of the integration sites, leaving the assay data the same.

For each assay, we examined the fold change of the original RPM against other cell types and compared with the fold change of the permuted RPM against other cell types. We did this for each 10 kbp window from the site of HPV integration up to 500 kbp away. We conducted a two-sided t-test on the differences (Figure 5a).

**Figure 5:**
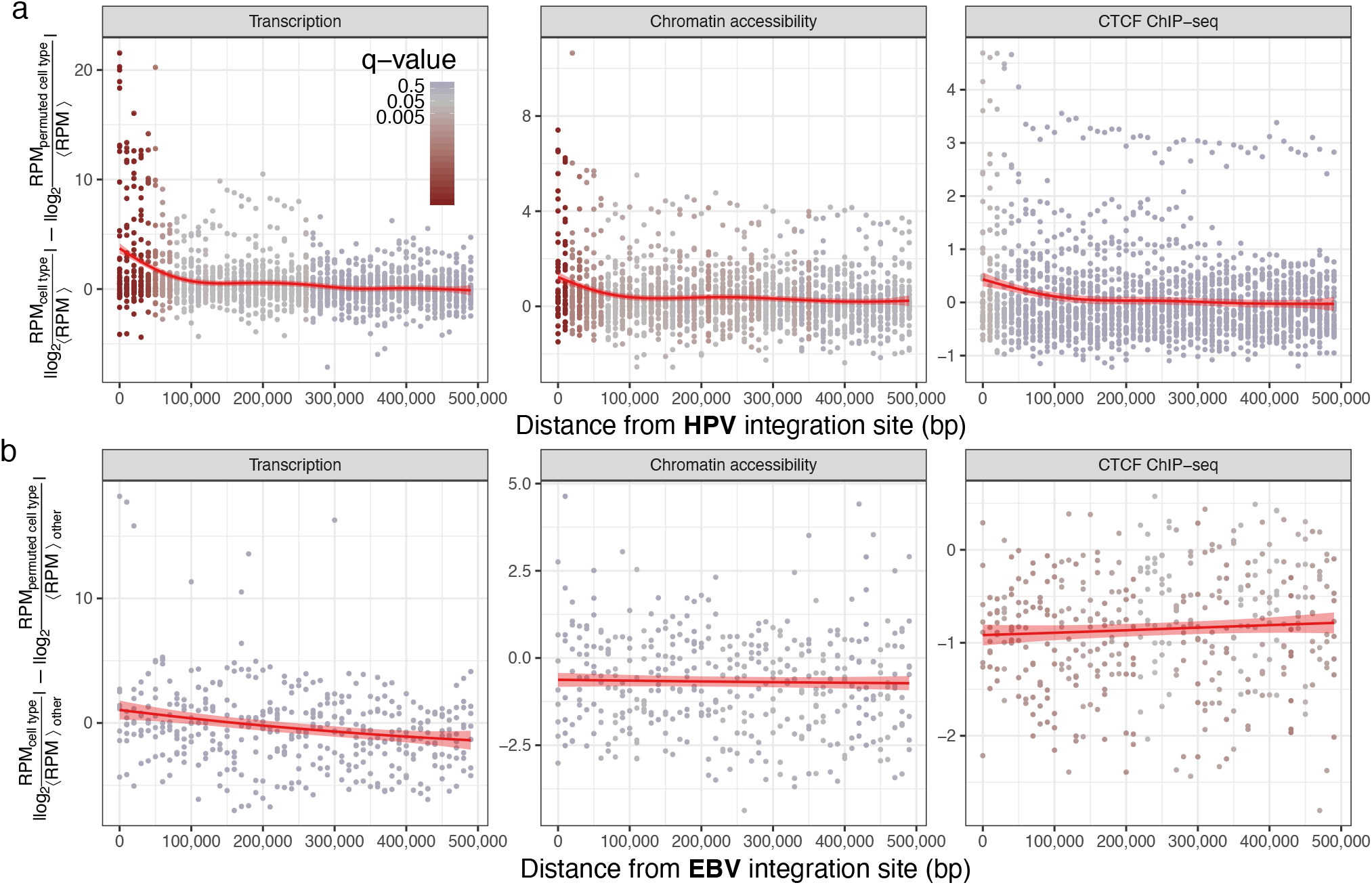
HPV integration dysregulated the epigenome and transcriptome up to 100 kbp away. **(a)** Difference in average RPM fold change in cell lines with HPV compared to 10 permuted controls. Each data point assesses the difference at a 10 kbp genomic bin. Color: q-value of t-test comparing the cell line with HPV integration to 10 permutations. Line: generalized additive model^32^ regression model on transcription data (RNA-seq; left), chromatin accessibility (ATAC-seq; middle), and CTCF presence (ChIP-seq; right). **(b)** Same as (a) but comparing GM12878 Epstein-Barr virus (EBV) integration sites to 2 other lymphoblastoid cell lines.

RNA-seq, ATAC-seq, and CTCF ChIP-seq significantly differed between the original and permuted measurements up to 100 kbp from the HPV integration sites (*q* < 0.05). HPV’s effect size on transcription, chromatin accessibility, and CTCF binding diminished as distance from the HPV integration sites increased (Figure 5a).

We hypothesized that changes in epigenome and transcriptome occurred due to a specific feature of the integrated HPV, and would not just arise from any genomic insertion. Under this hypothesis, we expected that the integration of the 170 kbp Epstein-Barr virus (EBV) would not induce similar changes to HPV. Therefore, we investigated how the transcriptome and epigenome changed at the EBV integration sites of 4 lymphoblastoid cell lines: GM12873, GM12878, GM23248, and GM23338 (Figure 5b; Supplementary Table 2).

Unlike with HPV, we detected no significant difference in transcriptome or epigenome within 100 kbp of EBV integration sites (Figure 5b). We observed more transcription around EBV integration sites, but no statistically significant difference after correcting for multiple comparisons (*q* > 0.37). GM12878 had less accessible chromatin and less CTCF binding compared to the other 2 lymphoblastoid cell lines when considering a larger region up to 500 kbp around EBV integration sites (*q* < 0.05). The magnitude of change, however, was relatively modest (RPM fold change of as much as −4) compared with the corresponding difference near HPV integration sites (RPM fold change of as much as 22).

#### 2.4.1 HPV integration dysregulates the local transcriptome of HPV^+^ carcinomas

Both cell lines derived from HNSC (93-VU147T and SCC-090) and cell lines derived from CESC (Caski and SiHa) displayed epigenomic and transcriptomic changes near HPV integration sites. To investigate how often outlier gene expression occurs due to biological variation other than HPV integration, we permuted RNA-seq data for these 4 cell lines, for TCGA HNSC samples, and for TCGA CESC samples. For HNSC and CESC datasets, we generated 100 permutations of samples as the background. For the 4 cell lines, however, we generated 10 permutations to avoid over-representation of the effects from the cell types with fewer viral integrations. In both the three original datasets and in corresponding permuted datasets each, we examined genes at thresholds *y* of expression fold change separated by intervals of 0.25. We identified those genes with expression exceeding *y* where the difference |TPMHPV − ⟨TPM_other_⟩| exceeded twice the standard deviation (SD) (Figure 6a).

**Figure 6:**
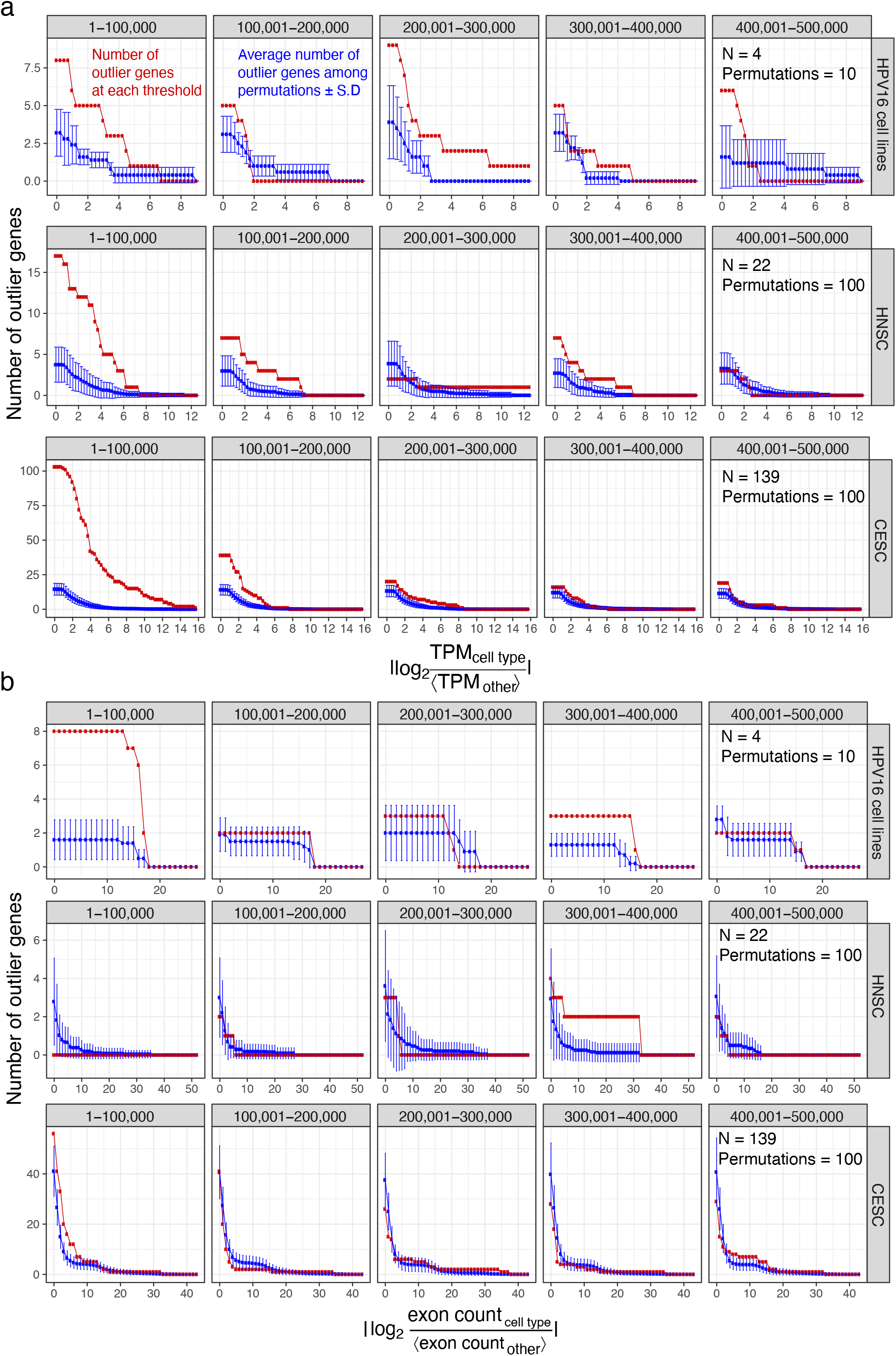
HPV integration dysregulated the epigenome and transcriptome up to 100 kbp away. **(a)** Complementary cumulative distribution function of number of outlier genes exceeding plotted absolute expression fold change ≥ horizontal axis values and |TPM_HPV_ − ⟨TPM_other_⟩| exceeding twice the SD. Top: 5 HPV16^+^ cell lines. Middle: HNSC patients. Bottom: CESC patients. Red: number of outlier genes in RNA-seq data; blue: mean number of outlier genes in 10 permutations of the samples; error bars: SD. **(b)** Similar to (a), but each showing the number of genes with absolute fold change in exon count > 1 and DEXSeq *q* < 0.2.

The original datasets contained more outlier genes passing a fold change cutoff of 2 compared to the permuted controls. The greatest deviation of the original datasets compared to the permuted datasets occurred within the 100 kbp window of HPV integration. In the 4 cell lines examined, we detected 8 outlier genes within 100 kbp of HPV integration, but a mean of 5 outlier genes in the 10 permuted datasets. Among HNSC tumours, we identified 16 outlier genes, far greater than the mean of 2 outlier genes in the permuted HNSC controls. We also identified 103 outlier genes among CESC tumours—as opposed to a mean of 13 outlier genes within the permuted CESC controls.

We performed a similar permutation analysis to investigate whether differential exon usage occurs due to biological variations other than HPV integration (Figure 6b). Within 100 kbp of HPV integration, we consistently identified more genes with differential exon usage in the original datasets compared to permuted controls. In the 5 cell lines examined, we found 8 genes with differential exon usage, but only a mean of 2 genes with differential exon usage among the permuted controls. In these 8 genes, absolute log_2_ exon count fold change (Equation 2) exceeded 13 (*q* < 0.2). We found similar results for CESC tumours.

We investigated whether the direction of the chimeric HPV transcript affects the magnitude of changes in neighbouring genomic regions. Although the most highly upregulated genes occurred near the HPV integration sites which induced the chimeric transcript downstream of the integration site, this represented a statistically insignificant difference (linear model *p* = 0.21).

### 2.5 HPV integration upregulates putative oncogenes

Having established that HPV integration results in changes in chromatin structure and dysregulated gene expression in cancer cell lines and patient tumours, we asked whether outlier expressed genes could play a driving role in tumourigenesis. We investigated the transcriptome of HPV^+^ HNSC and CESC tumours in TCGA. Out of the 71 HNSC HPV^+^ patients we examined, we found HPV integration sites in 22 of them by detecting transcribed chimeric sequences. Of these 22 patients, 16 (73%) displayed outlier expression of genes around HPV integration sites (Figure 7a). Among 228 CESC patients, 139 had transcribed chimeric sequences and therefore HPV integration sites. Of those 139 patients, 85 (61%) tumours displayed outlier expression of genes around HPV integration sites (Figure 7b).

**Figure 7:**
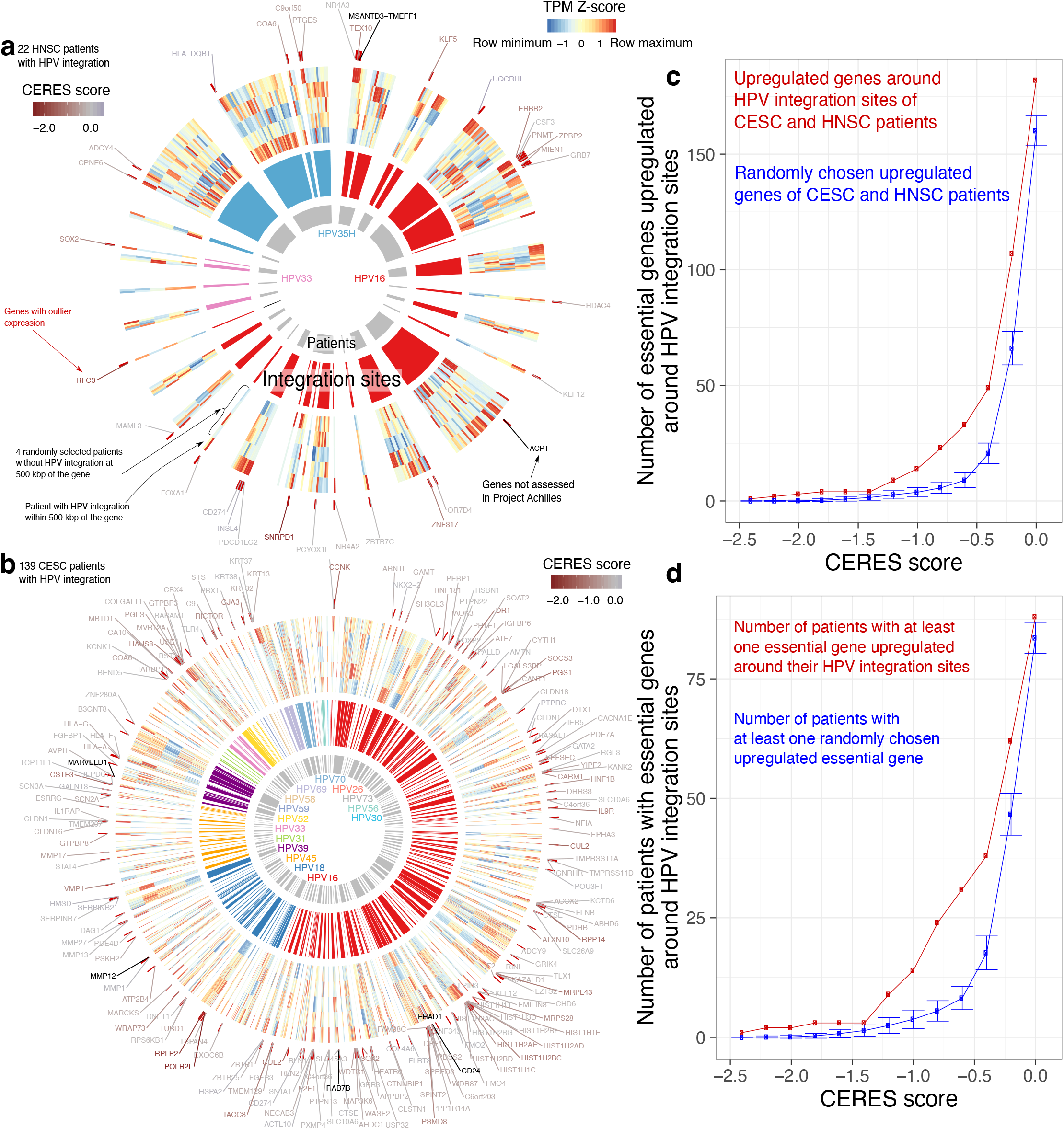
Outlier gene expression in HPV^+^ patients. **(a)** 26 TCGA HNSC patients with 3 strains integrated at 35 sites. Inner gray ring: each arc indicates a patient. Middle ring: individual HPV integration sites, with colour representing HPV strain. Outer ring: heatmap of expression of genes within 500 kbp of each integration site in the patient with HPV integration (peripheral) and 4 randomly selected patients without HPV integration around that gene (central). Red marks outside the heatmap: genes with outlier expression in the patient with HPV integration. Gene symbols: genes with outlier expression. Colour of the gene symbols indicate their CERES score. Red symbols indicate genes with negative CERES scores, essential to tumour viability. Black symbols indicate genes not assessed by Project Achilles. **(b)** 134 TCGA CESC patients with 12 strains integrated at 208 sites. **(c)** Number of genes upregulated (absolute log_2_ expression fold change > 1) in any of 160 patients from (a) and (b), where the gene’s CERES score < horizontal axis in at least one Project Achilles HPV^+^ cell line. Red: upregulated genes within 500 kbp of HPV integration sites. Blue: randomly chosen upregulated genes from the same patients. Blue data point: median of 100 permutations. Error bars: ±SD of 100 permutations. **(d)** Similar to (c), but instead the number of patients with at least one upregulated gene (absolute log expression fold change > 1), where the gene’s CERES score < horizontal axis in at least one Project Achilles HPV^+^ cell line.

Among the 5 cell lines, 26 HNSC tumours, and 85 CESC tumours, 231 genes in total showed outlier expression. HNSC patient TCGA-BA-5559, however, had an HPV integration at chr19:52,384,802, which disrupted the expression of 10 transcription factors with zinc finger domains (Figure 7a). Many genes with outlier expression around HPV integration sites, such as *FOXA1*^33^, *KLF12*^34^, *SOX2*^35^, *CUL2*^36^, *CD274*^37^, and *PBX1*^38^ have previously reported roles in tumourigenesis.

We used g:Profiler^39^ to identify dysregulated biological pathways more systematically (Supplementary Table 3). Genes with outlier expression around HPV integration sites enriched for the Gene Ontology (GO) terms “positive regulation of transcription by RNA polymerase II” (GO:0045944) and “cellular response to chemical stimulus” (GO:0070887, *q* < 0.05). Genes upregulated in HNSC patients enriched for the GO term “positive regulation of nucleobase-containing compound metabolic process” (GO:0045935). Genes upregulated in CESC patients enriched for “collagen catabolic process” (GO:0030574). These results suggest that genes upregulated upon HPV integration might activate transcription required for cellular proliferation, dysregulate cellular response to stress, anabolize nucleotides, and facilitate cellular invasion by degrading collagen within the basement membrane.

Project Achilles^40^ provides CRISPR-Cas9 screening data on the essentiality of 18,333 genes for the viability of 625 cancer cell lines. This includes 4 HPV^+^ cell lines (SiHa, Caski, SISO^41^, and SCC-152^27^). These datasets report a CERES score for each gene, which quantifies its essentiality for cancer proliferation and survival^40^. Non-essential genes have a median CERES score of 0 and common core essential genes have a median CERES score of −1.

Among the 223 upregulated genes around the integration sites of 101 HPV patient tumours, + 182 genes had negative CERES scores (Supplementary Table 4). For each patient, we performed 100 random permutations on the identity of the genes around their HPV integration sites, replacing them with other genes upregulated specifically in that patient (expression fold change > 2). Regardless of CERES score threshold used, we always found a higher number of upregulated essential genes in the original dataset than any of the permutation controls (Figure 7c). Also, more patient tumours had at least one upregulated essential gene around their HPV integration site, compared to randomly selected upregulated genes (Figure 7d).

## 3 Discussion

Several hypotheses can explain how HPV integration promotes tumourigenesis. Integration induces the expression of E6 and E7 either through disruption of the viral DNA-binding protein E2^42^, disruption of untranslated regions of E6 and E7^42^, or the creation of stable viral-host fusion transcripts^43^. Alternatively, certain integration sites may become genomically unstable, facilitating aberrant chromosomal rearrangements^26^ or may activate the expression of transposable elements, particularly short interspersed nuclear elements (SINEs)^44^. In many cases, transposable elements activate oncogenes and thereby initiate oncogenesis^45^. We propose that onco-exaptation of neighbouring genes by HPV could also prove sufficient to drive oncogenesis. Consistent with prior reports^9^,26,44,46, our results point to a separate mechanism whereby HPV integration leads to altered expression and splicing of neighbouring genes. Moreover, we identified active reorganization of local chromatin by CTCF binding to integrated HPV as a potential driver of local transcriptome dysregulation.

In agreement with previous reports^9^, our results show that HPV integration itself alters chromatin accessibility and the transcriptome in cell lines and patients. These changes may contribute to tumourigenesis by upregulating the expression of neighbouring genes, including some essential to tumour viability. In individual HPV integration sites, outlier expression of genes and changes in the epigenome occurred within 400 kbp of the integration. Examining integration sites in cell lines and patient tumours collectively uncovered significant chromatin, expression, and splicing differences within 100 kbp.

We identified a possible role for CTCF binding to integrated HPV in dysregulating the host chromatin and transcriptome. A conserved CTCF binding site distinguishes tumourigenic and non-tumourigenic HPVs^10^. In episomal HPV, knockout of this binding site enhances the expression of the E6 and E7 oncogenes^11^. A distinct role of the binding site in integrated HPV resolves this apparent paradox and explains its recurrence in tumourigenic strains.

Introduction of a new CTCF binding site by HPV may re-organize existing host topological domains. This can explain the extent of the changes in the chromatin and transcriptome seen here^12^. CTCF binding also plays a role in the life cycle of other DNA viruses, such as EBV, Kaposi’s sarcoma-associated herpesvirus, and herpes simplex virus 1 (HSV-1)^47^,48. We showed here, however, EBV integration does not lead to significant changes in chromatin at integration sites—only HPV integration does. Our data agree with previous work showing that only some changes to CTCF binding sites alter chromosome conformation^49^,50.

In a small number of observed cases, HPV integration has led to reorganization of topological domains^51^,52,53. These observations, however, have not linked the reorganization of the chromatin interactions to the conserved CTCF binding site of HPV as we propose here. Another important factor necessary for the formation of topological domain boundaries, SMC1, also binds the HPV genome at its CTCF binding sites^54^. Together with the existing literature on the epigenome of HPV, our results suggest that chromatin reorganization due to HPV integration occurs more frequently than previously appreciated.

We showed that HPV integration can increase the expression of neighbouring genes.We hypothesized that this, in turn, can predispose the host to tumour development. If true, the genomic position of the HPV integration site and the identity of its neighbouring genes should matter. Otherwise, we would expect HPV found in cancers integrated into genomic regions without any neighbouring oncogenes, since only a fraction of all genes can promote tumourigenesis. Reports on hotspot genomic regions in the host genome where HPV integrated^26^,44,55 and upregulated oncogenes around HPV integration sites^56^ support the hypothesis of increased local expression. The enrichment of HPV integration sites around genes^57^ and transposable elements, especially SINEs^44^, also supports this hypothesis.

If dysregulation of gene expression by HPV integration contributes to tumour development, we would expect to identify known oncogenes and master regulators of cancer-related pathways among the dysregulated genes in our analysis. Enrichment of these genes in growth-related pathways related to transcriptional regulation, nucleobase compound metabolism, and invasion-facilitating collagen catabolism, confirm our expectation. In agreement with recent studies^58^, upregulated genes around HPV integration sites enriched among the most essential genes compared to upregulated genes distant from HPV integration sites.

Most of the tumours we examined had chimeric transcripts that pinpointed integration sites. Only investigating these integration sites eliminated the possibility of detecting false positive integration sites. This approach, however, can miss some true integration sites that do not produce a chimeric transcript. It will also miss sites where one read of a pair maps completely to the virus and the other completely to the host. Future studies using long-read whole-genome sequencing or targeted approaches such as Tagmentation-assisted Multiplex PCR Enrichment sequencing (TaME-seq) could identify HPV integration sites more exhaustively^59^.

Regardless of these limitations, our results show that integration of HPV induces changes in local chromatin of the host and the local transcriptome. We predicted that these changes contribute to tumourigenesis. Our results suggest that interactions between integrated HPV chromatin and host chromatin triggers these changes, and that CTCF may play a key role in this process. Understanding the underlying mechanism of HPV–host chromatin interactions and their essentiality in tumourigenesis will better focus the future development of therapies for HPV^+^ cancers.

## 4 Methods

### 4.1 Multiple-testing correction

To control false discovery rate (FDR) over multiple comparisons, we used the Benjamini-Hochberg procedure^60^ to attain q-values^61^. We used q-value cutoff of 0.05 unless we indicated another threshold in the manuscript.

### 4.2 Genome assembly, annotations, and data processing

We generated a chimeric genome assembly and RefSeq gene transfer format (GTF) annotation of GRCh38 from Illumina iGenomes (https://support.illumina.com/sequencing/sequencing_software/igenome.html) and the National Center for Biotechnology Information (NCBI) RefSeq HPV16 K02718.1 assembly^62^. The resulting chimeric FASTA file had all the GRCh38 chromosomes, unplaced and unlocalized contigs, chrM (mitochondrial genome), EBV, and one additional chromosome containing the entire K02718.1 sequence. The GTF file contained all the Illumina iGenomes GRCh38 annotations and additional rows annotating K02718.1 coding sequences. For all experiments, we trimmed Illumina TruSeq adapters from FASTQ files with Trim Galore (version 0.4.4, https://www.bioinformatics.babraham.ac.uk/projects/trim_galore). For CTCF ChIP-seq, input control ChIP-seq, and ATAC-seq, we used Bowtie2^63^ (version 2.2.6) with default parameters to align FASTQ files to the chimeric GRCh38-HPV16 genome. For RNA-seq, we used STAR (version 2.6.0c)^64^, specifying options `--outFilterMultimapNmax 2 --genomeSAindexNbases 6 --alignSJoverhangMin 8 --alignSJDBoverhangMin 4 --outFilterMismatchNoverReadLmax 0.05` to align the FASTQ files to the chimeric GRCh38-HPV16 genome.

### 4.3 RNA-seq

#### 4.3.1 Library preparation and sequencing

We prepared samples for RNA-seq using the TruSeq Stranded Total RNA Sample Preparation kit with RiboZero Gold (Illumina, San Diego, CA). We performed RNA sequencing for each sample to ∼ 80 million paired-end 150 bp reads on an Illumina NextSeq 500 (Princess Margaret Genomics Centre, Toronto, ON). We collected input RNA using an AllPrep mini kit (Qiagen, Hilden, Germany).

#### 4.3.2 Bioinformatics analysis

We used StringTie^65^ (version 1.3.3b) to quantify TPM for genes in the chimeric GRCh38 annotation. We used DEXSeq (version 1.28.1) for alternative isoform analysis^66^. For DEXSeq, we downloaded Ensembl genes version 94 for compatibility with the DEXSeq protocol^67^. For each gene, we compared each sample against all the other samples. We repeated these steps for cell lines, HNSC, and CESC patients.

We generated a list of the exons with the most extreme difference in expression according to the DEXSeq negative binomial generalized linear model for all the genes around HPV integration sites. We quantified how the expression of each exon varies independent of the global expression of that gene (see subsubsection 4.3.2)^66^. For outlier exon expression, we again used a criterion of expression fold change > 2 compared to other cell lines:

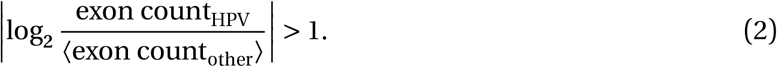

Instead of using the SD cutoff, we corrected the p-values for multiple testing using the Benjamini-Hochberg method^60^ and used a cutoff of *q* < 0.2 and minimum absolute fold change of 2 to select genes with alternative isoform expression.

#### 4.3.3 Identifying HPV-induced outlier expression

To determine whether HPV integration significantly changed gene expression, we examined changes in transcription of individual genes, as measured in TPM. We used two criteria to identify outlier changes in gene expression which occurred due to HPV integration. First, we calculated expression fold change dividing log_2_ TPM in the sample with HPV integrated at some locus (TPM_HPV_+) by median TPM in samples without HPV integrated at that locus (⟨TPM_other_⟩). For an HPV^+^ cell line, we only considered a gene an outlier if its expression fold change exceeded 2.

This meant a log_2_ fold change greater than 1:

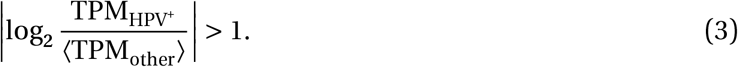

Fold change measurement, however, does not reflect dispersion in the expression of each gene. Second, therefore, we also required the difference in TPM to exceed at least twice the SD of TPM of that gene in other cell lines:

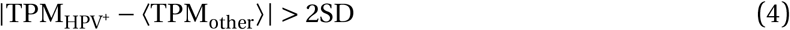

### 4.4 CTCF ChIP-seq

#### 4.4.1 Library preparation

We prepared 10 µL of both protein A and protein G beads through three washes of 5 mg/mL Dulbecco’s phosphate-buffered saline (dPBS) + bovine serum albumin (BSA). We added 10 µL’of polyclonal CTCF antibody (Cat No. 2899, Lot 002, Cell Signalling Technology, Danvers, MA; RRID:AB_2086794) to the beads in 300 µL dPBS + BSA and left it to bind for >6 h of rotation at 4 °C. After incubation, we washed the beads three more times with dPBS + BSA. Then, we resuspended the beads in protease inhibitor (PI) and 100 µL of modified radioimmunoprecipitation assay buffer (RIPA): 10 mmol/L Tris-HCl, pH 8.0; 1 mmol/L EDTA; 140 mmol/L NaCl; 1% volume fraction Triton X-100; 0.1% mass fraction SDS; 0.1% mass fraction sodium deoxycholate.

We trypsinized 1 million cells and then fixed for 10 min at room temperature in 300 µL of dPBS + 1% volume fraction formaldehyde. We added 15 µL of 2.5 mol/L glycine after fixing. Then, we washed the cells once in dPBS + PI before resuspending them in 300 µL of modified RIPA + PI. We sonicated the samples for 32 cycles of 30 s at full intensity using a Bioruptor Pico (Diagenode, Seraing, Belgium) and pelleted cell debris by spinning at 21,130 × g for 15 min. We set aside 15 µL of the supernatant as an input control, and diluted the remaining supernatant with 1700 µL of modified RIPA + PI and 100 µL of washed beads. We incubated the samples at 4 °C overnight with rotation. We washed the beads with the following cold buffers in order: modified RIPA, modified RIPA + 500 µmol/L NaCl, LiCl buffer (10 mmol/L Tris-HCl, pH 8.0; 1 mmol/L EDTA; 250 mmol/L LiCl; 0.5% mass fraction NP-40; 0.5% mass fraction sodium deoxycholate), and finally twice with TE buffer (10 mmol/L Tris-HCl, pH 8.0; 1 mmol/L EDTA, ph 8.0). We resuspended the samples and inputs in 100 µL of de-crosslinking buffer (1% volume fraction SDS, 0.1 mol/L NaHCO_3_) and incubated at 65 °C for 6 h. We cleaned the samples and inputs using the Monarch PCR & DNA clean-up kit (New England BioLabs, Ipswich, MA), prepared libraries using the ThruPLEX DNA-seq Kit (Rubicon Genomics, Ann Arbor, MI), and size selected to 240 bp–360 bp using a PippinHT 2% Agarose Cassette (Sage Science, Beverly, MA). For each sample, we sequenced three ChIP biological replicates and one input control to ∼25 million single-end 50 bp reads each on an Illumina HiSeq 2000 (Princess Margaret Genomics Core, Toronto, ON).

#### 4.4.2 Bioinformatics analysis

We used MACS2 (version 2.1.2) software^68^ to identify peaks and generate fragment pileup data using default parameters plus `--nomodel --bdg`, and using input as control. We also generated a log fold change enrichment bedGraph file by comparing fragment pileup to the input control lambda file generated by MACS2.

We used FASTQC^69^ (version 0.11.5) to assess the quality of ChIP-seq FASTQ files. After alignment with Bowtie2 and peak calling with MACS2, we used ChIPQC^70^ (version 1.18.2) to assess enrichment quality. Input controls always had less than 0.7% fraction of reads in peaks, while ChIP experiments had an average of 9.4% fraction of reads in peaks (SD 6.4%). We merged the three replicates and found the following number of peaks passing a threshold of 5% FDR and 5-fold enrichment over input control: 32,748 in 93-VU147T, 22,353 in Caski, 35,861 in HMS-001, 27,469 in SCC-090, and 37,161 in SiHa. Supplementary Tables 5–7 and supplementary files deposited in Zenodo (https://doi.org/10.5281/zenodo.3780364) provide additional quality control metrics.

### 4.5 ATAC-seq

#### 4.5.1 Library preparation and sequencing

We assessed open chromatin using OMNI-ATAC^71^ followed by size selection to 100 bp–600 bp using a PippinHT 2% Agarose Cassette (Sage Science, Beverly, MA) and paired-end 125 bp sequencing on an Illumina HiSeq 2500 to a depth of ∼60 million reads per sample (Princess Margaret Genomics Core, Toronto, ON).

#### 4.5.2 Bioinformatics analysis

We used MACS2 (version 2.1.2) software^68^ to identify peaks and generate fragment pileup data using default parameters and `--nomodel --shift -100 --extsize 200 --bdg --bampe`. For analysis of ATAC-seq peaks, we used an FDR threshold of 5%.

To visualize the chromatin accessibility signal of multiple samples at HPV integration sites, we used the FPM measurement of each sample divided to the maximum FPM of that sample in the chromosome of HPV integration. This ensured all of the values ranged between 0 and 1 in that chromosome.

### 4.6 TCGA datasets and analysis

#### 4.6.1 RNA-seq datasets

We downloaded GRCh37-aligned TCGA RNA-seq datasets for 295 CESC patients and 547 HNSC patients^30^. We extracted FASTQ files from the binary alignment map (BAM) files using bam2fastq (https://gslweb.discoveryls.com/information/software/bam2fastq). We aligned the samples back to the chimeric GRCh38-HPV16 genome using STAR^64^.

We used StringTie^65^ to quantify TPM for each of the experiments according to the chimeric GTF annotation of GRCh38 and HPV16. From the available 547 HNSC patients, we identified 58 as HPV^+^. We identified all of the 295 CESC patients as HPV^+^. We used DEXSeq for alternative isoform analysis^66^.

#### 4.6.2 ATAC-seq datasets

For the 9 TCGA HNSC patients with ATAC-seq data, we downloaded GRCh38-aligned BAM files. We extracted FASTQ files from the BAM files using bam2fastq (version 1.1.0, https://gslweb.discoveryls.com/information/software/bam2fastq), trimmed adapters and low-quality sequencing reads from the FASTQ files with Trim Galore, and aligned the samples back to the chimeric GRCh38-HPV16 genome using Bowtie2^63^ (version 2.2.6). We used MACS2 (version 2.1.2) software^68^ to identify peaks and generate fragment pileup data using default parameters and `--nomodel --shift -100 --extsize 200 --bdg --bampe`. For any analysis on ATAC-seq peaks, we used an FDR threshold of 5%.

### 4.7 EBV^+^ lymphoblastoid cell line datasets

We investigated how the transcriptome and epigenome changed at the EBV integration sites of 4 lymphoblastoid cell lines: GM12873, GM12878, GM23248, and GM23338. Of these cell lines, ENCODE supplies all 3 of total RNA-seq data, DNase-seq data, and CTCF ChIP-seq data for only GM12878 and GM23338. To provide 3 experiments for each assay, we added total RNA-seq and DNA-seq data from GM23248 and CTCF ChIP-seq from GM12873. For each of the 3 assays, this allowed us to compare potential differences arising from EBV integration in GM12878 to 2 other EBV^+^ lymphoblastoid cell lines.

### 4.8 Identifying HPV strains

For HNSC and CESC patients, we used the HPV strain reported previously^19^.

### 4.9 Identifying HPV integration sites

We developed Polyidus to identify HPV integration sites with chimeric sequencing reads from any paired-end sequencing data. First, Polyidus aligns reads to a viral genome. It allows for partial mapping using local alignment, and removes any sequencing fragment where neither read maps to the virus. Second, Polyidus aligns the selected reads to the host genome, permitting partial mapping. Third, Polyidus identifies *chimeric reads*: those reads mapped partially to the host genome and partially to the virus genome. Fourth, for each chimeric read, Polyidus reports the start and strand of integration in both the host and viral genomes. Polyidus also reports the number of chimeric reads supporting each integration site.

Polyidus finds highly confident integration sites which contain chimeric sequencing reads. Other methods perform the first two steps in reverse order^72^, resulting in slower performance. While some previous methods also align to the virus first^73^, either the software no longer appears available where specified at publication^74^,75, or they use BLAST^76^,77 instead of a faster short read aligner^78^. Unlike ViFi^44^, Polyidus requires that the chimera match an existing viral genome reference. Polyidus does not use non-chimeric fragments where one read maps entirely to host and one read maps entirely to virus genome.

Polyidus uses Bowtie2^63^ (version 2.2.6) and vastly speeds up integration site finding. Polyidus identified integration sites at an average of 8 core-hours on a 2.6 GHz Intel Xeon E5-2650 v2 processor and 4 GB of RAM for whole genome sequencing data. Previous methods^79^ require an average of 400 CPU core-hours.

We identified HPV integration sites in each sample using the sequence of the dominant HPV strain in that sample. We excluded any HPV integration site found in more than 1 patient to avoid overestimation of outliers at potential hotspots of frequent integration^26^. In some cases, we found more than one HPV integration site in a 20 kbp window in one patient. Since we used RNA-seq for identifying our integration sites, some of these integration sites might occur as a result of splicing between the integrated HPV and neighbouring host genomic regions. To avoid over-representing genomic regions with multiple integration sites, we only used the integration site with the highest number of chimeric sequencing reads.

## Availability

Polyidus provides a framework to catch chimeric sequences using Python. It is available on GitHub (https://github.com/hoffmangroup/polyidus) and deposited in Zenodo (https://doi.org/10.5281/zenodo.3780203). We deposited our datasets for RNA-seq, ATAC-seq, and CTCF ChIP-seq of 5 HPV^+^ cell lines in the Gene Expression Omnibus (GEO)^80^ (GEO accession: GSE143026) and other processed data in Zenodo (https://doi.org/10.5281/zenodo.3780364).

## Acknowledgments

We thank Jeff Bruce (ORCID: 0000-0002-6844-0286) for his assistance in data access. We thank Carl Virtanen (ORCID: 0000-0002-2174-846X) and Zhibin Lu (ORCID: 0000-0001-6281-1413) (University Health Network High-Performance Computing Centre and Bioinformatics Core) for technical assistance. We thank Elana Fertig (ORCID: 0000-0003-3204-342X) for providing histone modification ChIP-seq data for SCC-090. This work was supported by the Canadian Cancer Society (703827 to M.M.H.), Ontario Institute for Cancer Research Investigator Award (M.L.), the Princess Margaret Cancer Foundation with philanthropic support from the Joe and Cara Finley Centre for Head & Neck Cancer Research (S.V.B.) and the Gattuso-Slaight Personalized Cancer Medicine Fund (S.V.B. and M.M.H.), the Ontario Ministry of Training, Colleges and Universities (Ontario Graduate Scholarship to M.K.), the University of Toronto Faculty of Medicine Frank Fletcher Memorial Fund (M.K.), the Peterborough K.M. Hunter Charitable Foundation Graduate Award (M.K.), and the Parya Trillium Foundation Scholarship (M.K.).

## Author contributions

Conceptualization, M.K, M.L., S.V.B., and M.M.H.; Data curation, M.K.; Formal analysis, M.K.; Funding acquisition, M.L., S.V.B., and M.M.H.; Investigation, M.K., C.A., and A.R.; Methodology, M.K., C.A., S.V.B., M.L., and M.M.H.; Project administration, M.L., S.V.B., and M.M.H.; Resources, M.L., S.V.B., and M.M.H.; Software, M.K.; Supervision, M.L., S.V.B., and M.M.H.; Visualization, M.K.; Writing — original draft, M.K. and C.A.; Writing — review & editing, M.K., C.A., A.R., M.L., S.V.B., and M.M.H.

## Competing interests

The authors declare that they have no competing interests.

## Notes

https://github.com/hoffmangroup/polyidus

https://doi.org/10.5281/zenodo.3780203

https://doi.org/10.5281/zenodo.3780364

https://www.ncbi.nlm.nih.gov/geo/query/acc.cgi?acc=GSE143026

### Competing Interest Statement

The authors have declared no competing interest.

### Summary of Updates

Since HMS-001 lacks the genomic region of interested in the integrated form, we removed HMS-001 from all analyses and figures, except when it was used as a control for other cell types. Increased all the permutations from 10 to 100, when practical. Used SiHa CTCF ChIP-seq instead of TCGA-BA-A4IH chromatin accessibility in Figure 1a. Added chromatin accessibility peaks and summit locations to Figure 1b. Removed Figure 1c comparing host and viral CTCF binding, since it is addressed more systematically in Figure 3 and Figure 5. Replaced examples of HMS-001 with SiHa in Figure 2. Split previous Figure 5 into new Figure 5 and Figure 6. Added Supplementary Tables 5-7, which have quality control measurements for sequencing data.

